# Calsarcin-1 modulates the muscle LIM protein/Z-disc complex mechanosensing in familial hypertrophic cardiomyopathy

**DOI:** 10.64898/2026.11.15.766848

**Authors:** Xin Li, Yinsheng Lu, Xianmu Li, Muhammad Riaz, Thomas Blum, Hangqi Luo, Marie Hoareau, Yufeng Liu, Saba Fooladi, Qinzhe Xing, Ilhan Gokhan, Xia Li, Jinkyu Park, Lixia Yue, Stuart G. Campbell, Yibing Qyang

**Affiliations:** Yale Cardiovascular Research Center, Section of Cardiovascular Medicine, Department of Internal Medicine, Yale School of Medicine, New Haven, CT, USA; Yale Stem Cell Center, New Haven, CT, USA; Department of Pathology, Yale School of Medicine, New Haven, CT, USA; Department of Mechanical Engineering and Materials Science, Graduate School of Arts and Sciences, Yale University, New Haven, CT, USA; Department of Biomedical Engineering, Yale University, New Haven, CT, USA; Department of Cellular and Molecular Physiology, Yale School of Medicine, New Haven, CT, USA; Department of Physiology, College of Medicine, Hallym University, Hallymdaehak-gil, Chuncheon-si, Gangwon-Do, South Korea; Pat and Jim Calhoun Cardiology Center, University of Connecticut School of Medicine, Farmington, CT 06030, USA; Department of Cell Biology, University of Connecticut School of Medicine, Farmington, CT 06030, USA; Yale Biomedical Imaging Institute, Yale University, New Haven, CT, USA

**Author notes:** Xin Li, Yinsheng Lu, and Xianmu Li contributed equally. Correspondence (Y.Q.) and (S.G.C.).

**Keywords:** engineered heart tissue, EHT bioreactor, calsarcin-1, hypertrophic cardiomyopathy (HCM), muscle LIM protein (MLP)/Z-disc mechanosensory (MZM) complex

## Abstract

**BACKGROUND:** Hypertrophic cardiomyopathy (HCM) is an inherited cardiac disease mainly caused by sarcomeric mutations. The Z-disc is a critical mechanosensory interface linking sarcomeric stress to biochemical signaling, yet the role of calsarcin-1 in maintaining Z-disc integrity remains unclear. Patient-specific iPSC-CM-based engineered heart tissues (EHTs) are being developed for disease modeling and drug screening; however, the loading conditions to recapitulate pathological mechanical stress remain underdeveloped.

**METHODS:** HCM patient heart, three patient-specific iPSC lines (MLP-W4R;MYH7-R723C, MYH7-R663H, and MYBPC3-V321M), derived cardiomyocytes (iPSC-CMs), and engineered heart tissues (EHTs) were employed. Calsarcin-1 expression was assessed, and its role in MLP/Z-disc mechanosensing was investigated through gain- and loss-of-function approaches. An EHT bioreactor was developed to recapitulate HCM-associated elevated afterload, electrical pacing, and cyclic mechanical loading; isogenic wild type and MLP-W4R iPSC lines were generated by genome editing. Calsarcin-1 treatment to rescue the hypertrophic phenotype of isogenic (Iso) MLP-W4R iPSC-CM-EHTs was evaluated.

**RESULTS:** Calsarcin-1 was markedly downregulated in myocardium from an HCM proband carrying heterozygous MLP-W4R and MYH7-R723C variants, as well as in patient-specific iPSC-CMs and EHTs, accompanied by marked upregulation of the hypertrophic markers atrial natriuretic factor (*ANF*) and brain natriuretic peptide (*BNP*). Ectopic expression of calsarcin-1 via viral infection in MLP-W4R;MYH7-R723C iPSC-CMs stabilized the MLP/Z-disc machinery and reversed hypertrophy in all three patient-specific iPSC-CMs (MLP-W4R;MYH7-R723C, MYH7-R663H, and MYBPC3-V321M). Knockdown of calsarcin-1 attenuated the rescue effect of MLP overexpression in MLP-W4R;MYH7-R723C iPSC-CMs. Isogenic wild type and MLP-W4R iPSCs were generated by correcting mutations in MLP-W4R;MYH7-R723C iPSCs. Moreover, we developed a 3D-EHT bioreactor system that integrates controlled mechanical loading and electrical pacing to recapitulate the elevated afterload and cyclic mechanical stress characteristic of HCM. The additional mechanical resistance imposed by high afterload prolongs TTP and RT50 and increases the normalized force-time integral (nFTI), suggesting that excessively high afterload drives pathological remodeling in IsoMLP-W4R-iPSC-CM-EHTs. More importantly, calsarcin-1 significantly rescued both systolic and diastolic dysfunction of IsoMLP-W4R-iPSC-CM-EHTs exposed to pathological high afterload.

**CONCLUSIONS:** Calsarcin-1 is a nodal regulator of pathological cardiac remodeling that restores MLP/Z-disc mechanosensing. A patient-derived EHT bioreactor model recapitulating pathological HCM loading identifies calsarcin-1 as a potential therapeutic target.

## INTRODUCTION

Hypertrophic cardiomyopathy (HCM), a hereditary disorder of cardiac muscle, affects approximately 1 in 500 individuals and contributes to sudden cardiac death in adolescents and young adults^1^. HCM is often caused by pathogenic variants in genes encoding sarcomeric proteins, such as β-myosin heavy chain (MYH7) and myosin-binding protein C (MYBPC3)^2^. β-blockers, calcium-channel blockers, and recently developed cardiac myosin inhibitors are used clinically to attenuate contractile hyperfunction and downstream hypertrophic remodeling. Yet more than three decades after the first sarcomeric mutation was identified, the molecular events linking hypercontractile sarcomeric variants to cardiomyocyte hypertrophy and disarray remain incompletely understood^3^.

Muscle LIM protein (MLP) couples sarcomere-generated mechanical force to signaling cascades that govern the hypertrophic response, and the integrity of this MLP-dependent Z-disc mechanosensing (MZM) machinery is required for physiological mechanotransduction in the mammalian heart^4,5^. We previously demonstrated that the MLP-W4R variant identified in familial HCM is susceptible to degradation in the hypercontractile MYH7-R723C genetic background, leading to destabilization of this MZM machinery and consequent aberrant activation of calcineurin–NFAT signaling and pathological hypertrophy^6^. This establishes the critical role of the MLP-dependent mechanosensing machinery in linking sarcomere dysfunction to disease. However, the molecular determinants that modulate the structural and signaling integrity of the MLP-dependent mechanosensing machinery during disease progression remain incompletely defined.

Calsarcin-1 (encoded by MYOZ2) is a sarcomeric Z-disc protein that negatively regulates calcineurin signaling by anchoring calcineurin to the Z-discs of the sarcomere, thereby preventing excessive activation of the calcineurin pathway and modulating the hypertrophic events in response to biomechanical stress in murine studies^7,8^. However, the role of calsarcin-1 as an essential molecular determinant that preserves the integrity of the MLP-dependent Z-disc mechanosensing machinery in human familial HCM remains unknown.

Herein, we showed that calsarcin-1 expression was markedly reduced in myocardium from the HCM patient heart carrying the MLP-W4R;MYH-R723C mutations as well as in patient-specific induced pluripotent stem cell–derived cardiomyocytes (iPSC-CMs) and engineered heart tissues (EHTs). Restoring calsarcin-1 expression stabilized the MLP-dependent Z-disc mechanosensing machinery, reversed pathological hypertrophy, and attenuated abnormal contractile and relaxation function in isogenic MLP-W4R iPSC-CM-EHTs. These findings identify calsarcin-1 as a previously unrecognized molecular determinant that preserves the integrity of the MLP-dependent Z-disc mechanosensing machinery, thereby establishing the foundation for a novel therapeutic strategy for hypertrophic cardiomyopathy.

## METHODS

### Cell lines and culture condition

Human induced pluripotent stem cells (iPSCs) were maintained on mouse embryonic fibroblast (MEF) feeder layers in human embryonic stem cell (hESC) medium. Cells were cultured until colonies reached approximately 70%–80% confluence. For passaging, iPSC colonies were treated with Dispase II (1 U/mL; STEMCELL Technologies, 07923) for 8–9 min and gently detached using a 1-mL pipette tip. The detached colonies were transferred to a 15-mL conical tube and centrifuged at 200 rpm for 4 min to minimize MEF contamination. After removing the supernatant, iPSC colonies were resuspended in hESC medium supplemented with 10 μM Y-27632 and transferred onto freshly pre-seeded MEF feeder layers.

HEK293T cells were maintained in DMEM supplemented with 10% fetal bovine serum (FBS) and 1% penicillin-streptomycin and cultured at 37°C in a humidified atmosphere containing 5% CO₂. For lentiviral production, HEK293T cells were seeded at an appropriate density and transfected with the lentiviral transfer plasmids encoding MLP or Calsarcin-1 for overexpression, or shRNA targeting the indicated gene or scramble shRNA as a control, together with the packaging plasmid psPAX2 and the envelope plasmid pMD2.G, using polyethylenimine (PEI). Culture medium was replaced 6 hrs. after transfection. Viral-containing supernatants were collected at 48 and 72 hrs. after transfection, clarified by centrifugation, and used for subsequent transduction experiments.

### Mouse strains and husbandry conditions

CF1 mice were obtained from Charles River Laboratories and maintained under standard laboratory conditions in a temperature- and humidity-controlled animal facility with a 12-h light/12-h dark cycle and ad libitum access to food and water. Embryos at embryonic day (E) 12–13 were collected from timed-pregnant CF1 females and used for the isolation of mouse embryonic fibroblasts (MEFs). All animal procedures were performed in accordance with institutional guidelines and were approved by the Institutional Animal Care and Use Committee (IACUC) at Yale University.

### Immunofluorescence Staining

Paraffin-embedded human heart sections from healthy donors and patients with hypertrophic cardiomyopathy (HCM; MLP-W4R and MYH7-R723C) were deparaffinized in xylene for 10 min and rehydrated through a graded ethanol series (100%, 95%, 95%, and 75%; 5 min each). Sections were washed with PBS and subjected to heat-induced antigen retrieval in 1× Unmasking Solution at 97–100°C for 20 min. hiPSC-derived cardiomyocytes (hiPSC-CMs) were fixed with 4% paraformaldehyde for 15 min at room temperature and washed with PBS. Heart sections and hiPSC-CMs were then permeabilized with 0.1% Triton X-100 in PBS for 20 min and blocked with 5% BSA or goat serum in PBS for 30 min at room temperature. Samples were incubated overnight at 4°C with primary antibodies against calsarcin-1 (Abcam, ab237053) and muscle LIM protein (MLP; Santa Cruz Biotechnology, sc-166930) for heart sections, or mouse monoclonal anti-cardiac troponin T (cTnT) antibody [1C11] (Abcam, ab8295) for hiPSC-CMs. Following three washes with 0.1% Tween-20 in PBS, samples were incubated for 1 h at room temperature with the appropriate secondary antibody: goat anti-rabbit IgG Alexa Fluor 488 (1:500; Invitrogen, A-11008) or goat anti-mouse IgG Alexa Fluor 647 (1:500; Invitrogen, A-21235). Nuclei were counterstained with DAPI Fluoromount-Gold mounting medium (SouthernBiotech, 0100-20). Images were acquired using a Leica TCS SP8 confocal microscope (Leica Microsystems).

### Mouse embryonic fibroblast (MEF) isolation

Mouse embryonic fibroblasts (MEFs) were isolated from E12–13 CF1 mouse (Charles River Laboratories) embryos. Following removal of the head, visceral organs, and major blood vessels, embryos were minced and digested with 0.05% Trypsin-EDTA (Gibco, 25-300-054) to generate a single-cell suspension. MEFs were cultured in MEF medium consisting of DMEM (Gibco, 11965-092) supplemented with 10% fetal bovine serum (FBS; Gibco, 10082-147), 1% non-essential amino acids (Gibco, 11140050), 1% Penicillin–Streptomycin (Gibco, 15070063), and 2 mM L-glutamine (Gibco, 25030149). MEFs were expanded to passages 2–3 and treated with Mitomycin C (Selleckchem; S8146) to induce mitotic arrest prior to use as feeder layers for iPSC maintenance. All animal procedures were approved by the Institutional Animal Care and Use Committee (IACUC) of Yale School of Medicine and conducted in accordance with the NIH Guide for the Care and Use of Laboratory Animals.

### Human induced pluripotent stem cells cardiac differentiation

Cardiac differentiation was performed as previously described^6,35^. Briefly, for cardiac differentiation, human iPSC colonies were passage from MEF feeder layers using Dispase II (1 U/mL; STEMCELL Technologies, 07923), collected as small aggregates, and resuspended in in mTeSR™ Plus medium (Stemcell Technologies, 100-0276) supplemented with 5 μM ROCK inhibitor Y-27632 (Tocris Bioscience, 1244) and replated onto Growth Factor Reduced (GFR) Matrigel-coated plates (Corning; 1:60 dilution in DMEM/F12) at a 1:1.5 split ratio. Differentiation was initiated once cultures reached 80–95% confluency, typically within 2–3 days.

On day 0, cells were treated with 20 μM CHIR99021 (Selleckchem, S2924) in a differentiation medium consisting of three parts RPMI-1640 supplemented with B27 minus insulin (Gibco, A18956-01) and one-part mTeSR™ Plus medium. Twenty-four hours later, replace the medium with RPMI/B27 minus insulin. On day 3, cells were treated with the Wnt inhibitor IWP-4 (5 μM; Stemolecule, 04-0036) in RPMI/B27 minus insulin, followed by a medium change on day 5. Thereafter, the medium was refreshed every other day. Spontaneously beating cardiomyocytes generally appeared between days 9 and 11. On day 9, cultures were transitioned to RPMI-1640 supplemented with B27 containing insulin (Gibco, 17504044).

To enrich the cardiomyocyte population, cells underwent metabolic selection beginning on day 12 using glucose-free DMEM (Gibco, 11966025) supplemented with 4 mM sodium lactate (Sigma-Aldrich, L7022) for 4 days. Following selection, cells were returned to RPMI-1640 supplemented with B27 and insulin and allowed to recover for an additional 2 days. Purified cardiomyocytes were harvested on day 18 and used to generate engineered heart tissue (EHT).

### Engineered heart tissue (EHT) generation

Engineered heart tissues (EHTs) were fabricated using induced pluripotent stem cell-derived cardiomyocytes (iPSC-CMs) seeded into decellularized porcine myocardial scaffolds as previously described^12^. Porcine myocardium was sectioned into 150-µm slices, patterned by laser cutting, and decellularized through sequential treatment with cell lysis buffer (10 mM Tris, 0.1% w/v EDTA, pH 7.4) for 2.5 h and 0.5% (w/v) sodium dodecyl sulfate (SDS) in DPBS for 60 min under gentle agitation at room temperature. Following three washes in DPBS, scaffolds were conditioned overnight in DMEM supplemented with 10% fetal bovine serum and 1× Penicillin–Streptomycin.

On day 14 of differentiation, iPSC-CMs were dissociated with TrypLE (Gibco, 12605010) at 37°C for 10–15 min. Quench enzymatic activity by dilution in MEF medium, then collect cells by centrifugation (1,000 rpm, 5 min). The cell pellet was resuspended in MEF medium containing 5 μM Y-27632. For each EHT, 1.35 × 10^6^ iPSC-CMs were combined with 0.15 × 10^6^ human cardiac fibroblasts (Promocell, C-12375) and seeded into a preconditioned myocardial scaffold. EHTs were maintained under standard culture conditions, with medium changed to DMEM supplemented with 1× B-27 Supplement with insulin (Gibco, 17504044) and 1× Penicillin–Streptomycin (Gibco, 15070063) beginning 48 h after seeding. EHTs were subsequently cultured in this medium until functional assessment.

### Bioreactor design and fabrication

This device can apply iPSC-CM-EHTs to experience precisely controlled afterloads by applying either shortening or static culture in a single dimension. Programming the device to follow a predefined trajectory synchronized with electrical pacing ensures precise, reproducible mechanical stimulation of the EHTs. A typical motion profile for simulating a low afterload condition involves a 10% shortening regime at 1 Hz. The motion is defined over a 1000 ms cycle, discretized into 48 time points [‘t = linspace(0,1000,48)’] to achieve smooth and controlled mechanical deformation. The position profile is designed to begin and return to zero, mimicking the natural contraction-relaxation cycle of EHTs. Electrical pacing is applied at 135 ms within the cycle, strategically timed to synchronize with the contraction-relaxation dynamics while avoiding the first and last 10 ms to minimize motion artifacts and ensure accurate force recordings. This configuration facilitates precise afterload application along one dimension, enabling reproducible and physiologically relevant contraction responses under controlled mechanical conditions. In contrast to dynamic EHT training under low afterload conditions, static training involves maintaining the EHTs in a fixed position without mechanical movement while applying electrical pacing at 1 Hz. This approach provides EHTs with sustained tension, mimicking high afterload conditions during culture.

A custom dynamic bioreactor was developed to apply programmable mechanical stimulation to EHTs under controlled afterload conditions. The system integrates a voice coil linear actuator (H2W VCS03-050-BS-01-C) mounted on a precision linear rail, enabling smooth uniaxial motion with high temporal resolution and minimal friction. Displacement is monitored by an LM10 incremental magnetic encoder (Renishaw RLS), which provides non-contact, high-resolution position feedback suitable for incubator environments. The encoder’s magnetic scale is attached to the moving platform, while the readhead remains fixed to the chassis, enabling accurate real-time tracking of actuator position.

Motion is controlled by a Galil DMC-4103 controller and D3540 servo amplifier, executing pre-programmed trajectories via closed-loop PID control. Motion profiles, such as a 10% sinusoidal shortening at 1 Hz (48 points over 1000 ms), are uploaded via GalilTools and synchronized with electrical pacing (e.g., a 135 ms delay) to achieve physiologically relevant contraction cycles. The dynamic bioreactor chamber is fabricated from autoclavable materials and designed for sterile operation within a standard incubator. Electrical and signal cables are routed through sealed ports, and the modular assembly facilitates cleaning and reusability. The platform supports synchronized electromechanical conditioning, enabling controlled, reproducible stimulation of cardiac tissues. For the high-afterload group, the EHTs are placed in a 6-well plate cell culture dish and assembled with a carbon electrode pacing board.

### Engineered Heart Tissue Functional Assessment

The contractile function of engineered heart tissues (EHTs) was evaluated using a custom muscle mechanics platform, as previously described^12^. Following removal from culture, EHTs were mounted between a force transducer and a motorized length controller and submerged in a temperature-regulated organ bath. Tissues were continuously superfused with oxygenated Tyrode’s solution containing 140 mM NaCl, 5.4 mM KCl, 1 mM MgCl₂, 25 mM HEPES, 10 mM glucose, and 1.8 mM CaCl₂ (pH 7.35) and electrically stimulated via field electrodes at 1 Hz (100 mA, 10 ms pulse duration). All recordings were performed at 36°C. After equilibration and mechanical preconditioning, EHTs were stretched to 110% of their resting length (10 mm) for functional measurements. Force recordings from 20 consecutive contractions were averaged for each construct. Peak twitch force, time to peak force (TTP), and time to 50% relaxation (RT50) were subsequently determined as indices of systolic and diastolic performance.

### CRISPR/Cas9-Mediated Gene Correction

Correction of the MYH7-R723C variant was achieved using a CRISPR/Cas9-mediated homology-directed repair (HDR) strategy. A single-guide RNA (sgRNA) was designed using an online CRISPR design platform (https://chopchop.cbu.uib.no/) and cloned into a Streptococcus pyogenes Cas9 expression vector. Correct insertion of the sgRNA sequence was verified by Sanger sequencing. To serve as the HDR donor, a 901-bp repair template (Quintara Bioscience) containing the wild-type sequence was cloned into pUC19. In addition to correcting the pathogenic variant, the donor construct incorporated silent nucleotide substitutions that introduced a BamHI restriction site for clone screening and disrupted the PAM sequence to minimize Cas9 re-cutting following repair.

Human iPSCs maintained on Growth Factor-Reduced (GFR) Matrigel were co-transfected with the sgRNA-Cas9 plasmid, donor template, and a GFP reporter plasmid using Lipofectamine™ LTX. Twenty-four hours after transfection, the culture medium was replaced, and cells were maintained for an additional 48 hours before enrichment of GFP-positive cells by fluorescence-activated cell sorting (FACS). Sorted cells were replated onto GFR-Matrigel-coated culture dishes and expanded under hypoxic conditions (5% O₂). Individual colonies were manually picked after approximately 2 weeks and further expanded for genotyping. Targeted integration was assessed by PCR amplification of a 701-bp genomic fragment encompassing the edited locus, followed by BamHI restriction analysis. Candidate HDR-positive clones were verified by Sanger sequencing.

### TALEN-Mediated Correction of the MLP-W4R Variant

Targeted correction of the heterozygous MLP-W4R mutation was performed using a Transcription Activator-Like Effector Nuclease (TALEN)-based genome editing approach. TALEN pairs spanning the mutant locus were synthesized (Invitrogen) and assembled into a Gateway-compatible backbone before subcloning into a pEF1α-driven expression vector optimized for use in human iPSCs. A 135-bp antisense single-stranded oligodeoxynucleotide (ssODN) carrying the wild-type sequence served as the repair template. To facilitate the identification of homologous recombination events, the donor template included a silent nucleotide substitution that generated a diagnostic MfeI restriction site.

Human iPSCs maintained on Geltrex-coated plates in Essential 8™ medium were transfected at 40–50% confluency using Lipofectamine LTX and Plus reagents. The transfection mixture contained plasmids encoding the left and right TALEN arms, the ssODN repair template, and a GFP reporter construct. A parallel GFP-negative control was included to establish fluorescence gates for cell sorting. Cells were pretreated with ROCK inhibitor prior to transfection and returned to fresh culture medium following complex uptake. After 48 hours, GFP-positive cells were isolated by fluorescence-activated cell sorting (FACS) and replated at low density (5,000–10,000 cells per dish) onto Geltrex-coated culture dishes. Cultures were maintained under hypoxic conditions (5% O₂) to support clonal expansion. Emerging colonies were manually isolated after approximately 10 days and screened for targeted repair. Clones with successful correction of the MLP-W4R allele were identified by genotyping and restriction analysis, and the MYH-R723C;MLP-W4R corrected lines were selected for karyotype assessment and subsequent cardiac differentiation studies.

### Western blotting

Total protein lysates were extracted from day 35 hiPSC-CMs or engineered heart tissues (EHTs) using RIPA lysis buffer (Millipore, 20-188) supplemented with cOmplete™ Protease Inhibitor Cocktail (Roche; C756U27) and Phosphatase Inhibitor Cocktail 2 (Sigma-Aldrich; P5726), followed by sonication. Protein concentration was quantified, and 30–50 μg of total protein was resolved on an SDS-PAGE gel and transferred onto polyvinylidene fluoride (PVDF) membranes pre-activated with methanol. Membranes were blocked with 5% milk, incubated with specific primary antibodies, and then incubated with horseradish peroxidase (HRP)-conjugated secondary antibodies. Detection was performed using SuperSignal™ West Pico PLUS Chemiluminescent Substrate (ThermoFisher Scientific; 34579), and protein bands were visualized using the GBOX Chemi XX6 Gel Documentation and ECL Detection System. Primary antibody: rabbit anti-MLP (Abcam; ab42504), anti-MYOZ2 (MYBioSource; MBS4505333), anti-Gapdh (Abcam; ab8245); HRP-conjugated secondary antibody: horse anti-mouse IgG (Vector Laboratories; PI-2000), horse anti-rabbit IgG (Vector Laboratories; BA-1100-1.5).

#### Real-time qPCR

Total RNA was isolated from cells or tissues using TRIzol reagent (Invitrogen) according to the manufacturer’s instructions. RNA concentration and purity were determined spectrophotometrically. Complementary DNA (cDNA) was synthesized from 0.5 μg of total RNA using the iScript™ cDNA Synthesis Kit (Bio-Rad; 1708890) following the manufacturer’s protocol. Quantitative PCR was performed using iTaq Universal SYBR Green Supermix (Bio-Rad; 1725120) on a CFX Opus Real-Time PCR System (Bio-Rad). Reactions were carried out in triplicate under standard cycling conditions. Gene-specific primers were designed and commercially synthesized. Relative gene expression was calculated using the comparative Ct (2−ΔΔCt) method, with GAPDH serving as the internal reference gene. Data are presented as fold change relative to the control group.

### Quantification and statistical analysis

Data quantification and statistical analyses were performed using GraphPad Prism 9. All data are presented as mean ± SEM (standard error of the mean). Normality was evaluated using the Shapiro–Wilk test. For two-group comparisons, the two-tailed unpaired Student’s t test was used when data were normally distributed, whereas the two-tailed unpaired Mann–Whitney U test was used when data were not normally distributed. Comparisons among multiple groups were performed using one-way ANOVA followed by Tukey’s multiple comparisons test. For analyses involving two independent variables, two-way ANOVA with Tukey’s multiple comparisons test was used. A *P* value <0.05 was considered statistically significant.

### Antibodies and reagents

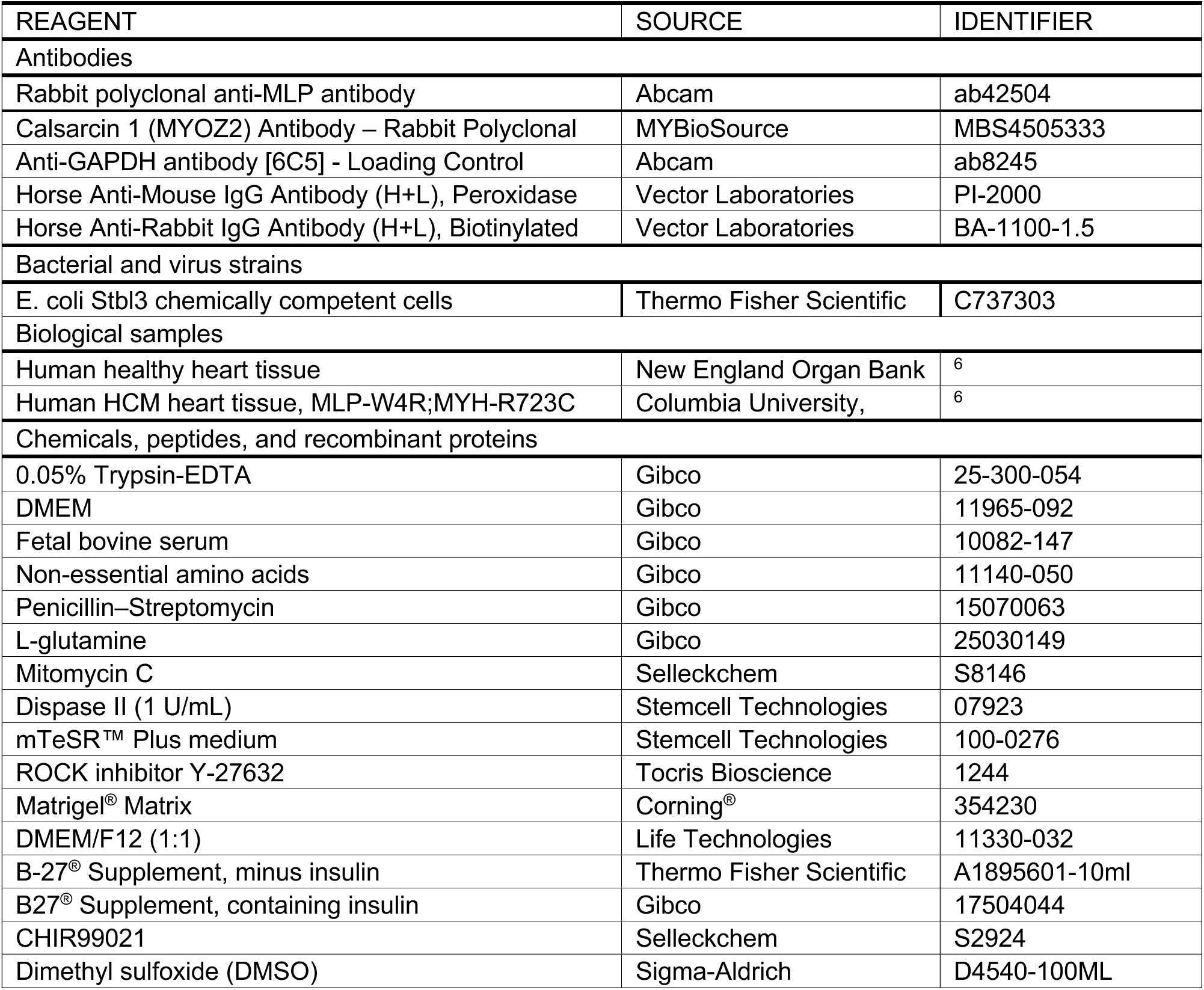
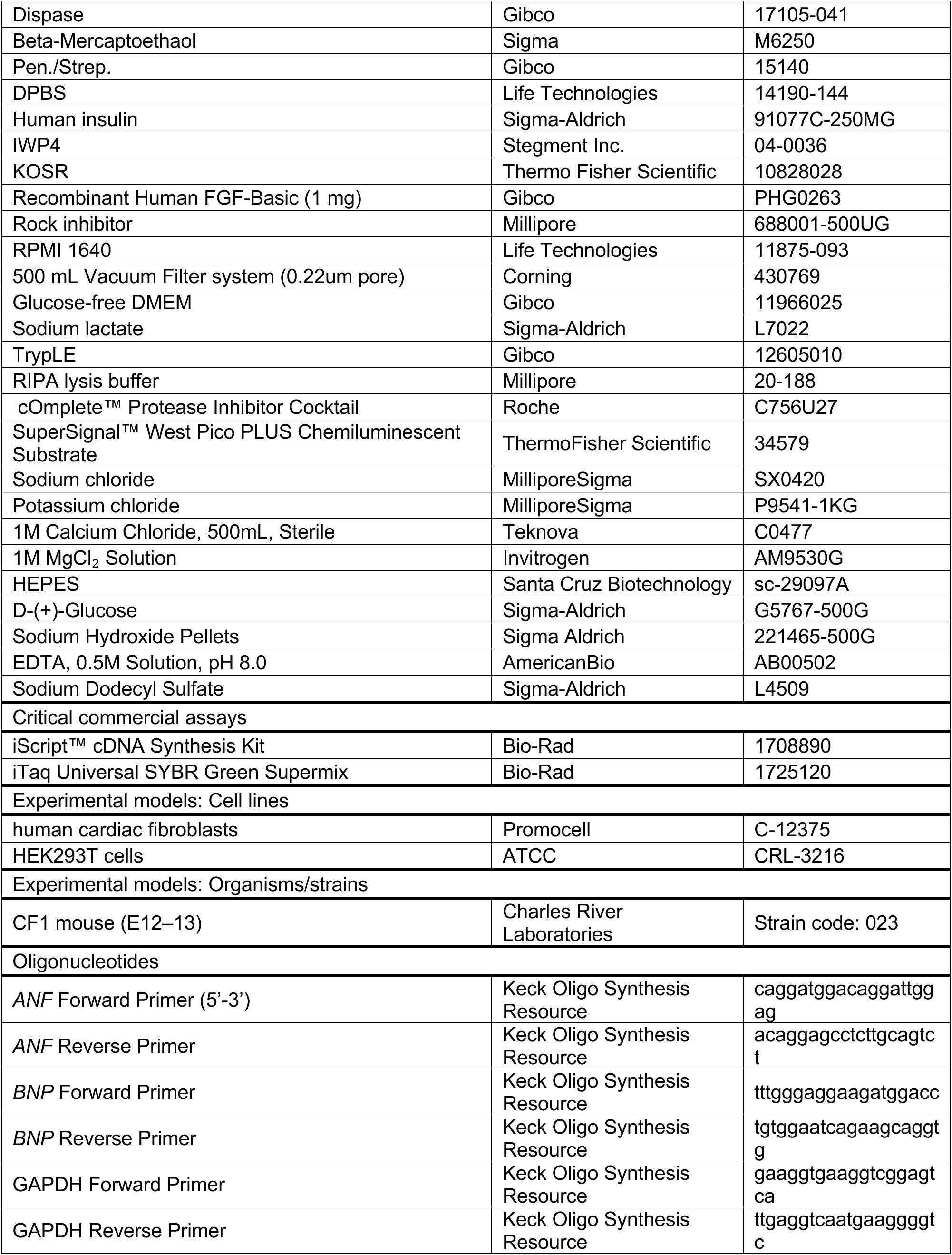
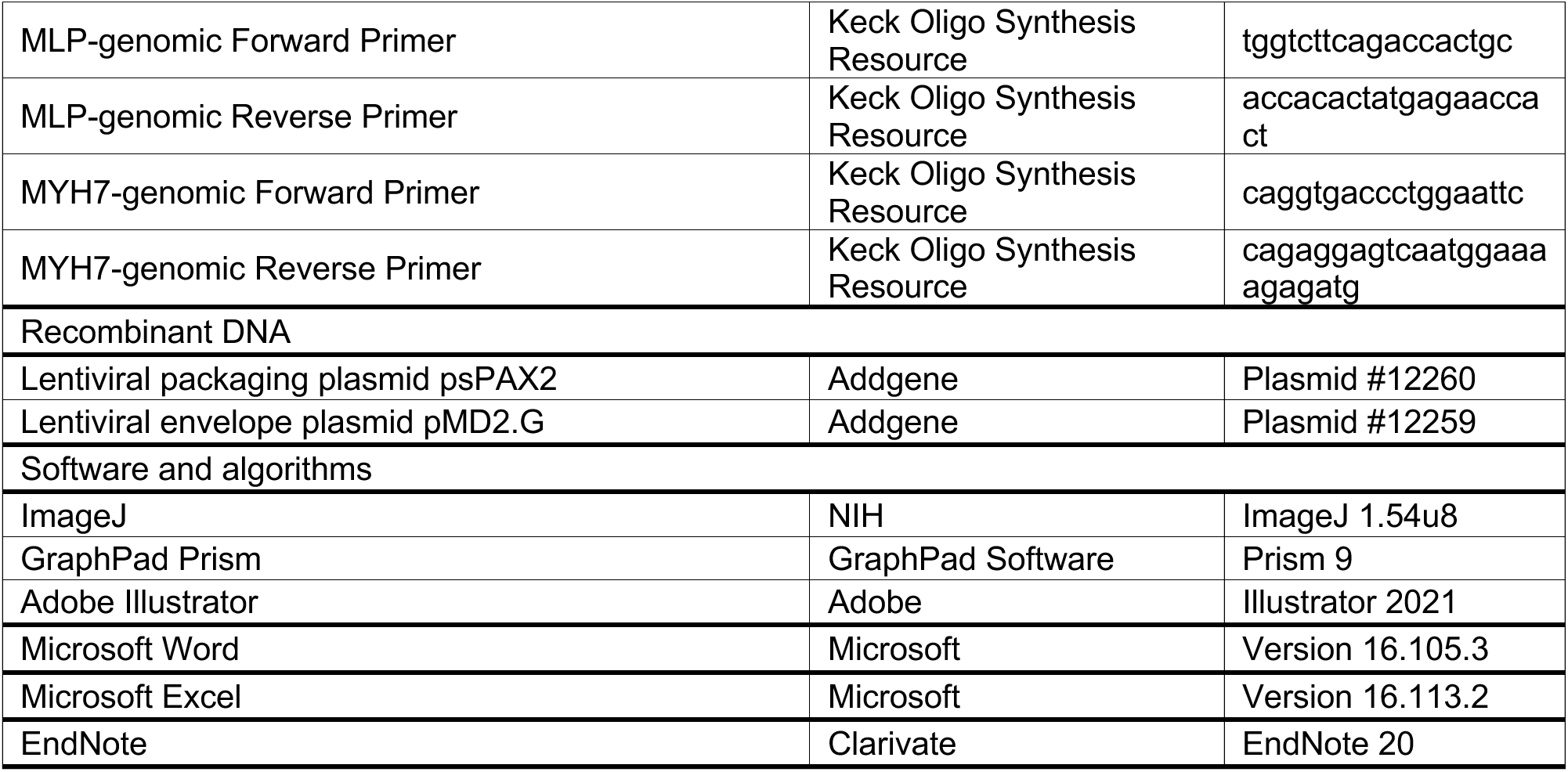

## RESULTS

### Downregulation of calsarcin-1 in the familial hypertrophic cardiomyopathy patient’s heart and iPSC-derived cardiomyocytes

We first explored the differential calsarcin-1 expression in the HCM proband’s heart, which carries two heterozygous mutations, MLP-W4R and MYH7-R723C (**Figure 1A**). While immunofluorescence staining showed regular, well-defined striated patterns with organized myofilaments in the healthy control (CTL) donor heart, the striated patterns were largely disrupted in the MLP-W4R;MYH7-R723C HCM patient heart. Intriguingly, the expression of calsarcin-1 appeared to be reduced in the MLP-W4R;MYH7-R723C proband myocardium compared to that in the young adult healthy heart (CTL, **Figure 1B**). Induced pluripotent stem cells (iPSCs) from the MLP-W4R;MYH7-R723C proband and healthy control donor (CTL) were differentiated into beating cardiomyocytes (iPSC-CMs) to determine the role of calsarcin-1 in human HCM. The gene expression levels of HCM-associated markers, including atrial natriuretic factor (*ANF*) and brain natriuretic peptide (*BNP*)^9–11^, were higher in the MLP-W4R;MYH7-R723C iPSC-CMs (**Figure 1C**). Meanwhile, calsarcin-1 protein expression in MLP-W4R;MYH7-R723C proband iPSC-CMs was significantly lower compared with that in CTL iPSC-CMs (**Figure 1D**). Notably, MLP protein levels also declined, confirming our previous finding of susceptibility to MLP degradation in MLP-W4R;MYH7-R723C iPSC-CMs^6^. Moreover, engineered heart tissues (EHTs) were generated by seeding CTL or MLP-W4R;MYH7-R723C iPSC-CMs with human cardiac fibroblasts (HCF) into a laser-cut decellularized porcine cardiac extracellular matrix (ECM) scaffold^12^. The expression level of calsarcin-1 protein in MLP-W4R;MYH7-R723C iPSC-CM-EHTs was significantly lower compared to that in CTL iPSC-CM-EHTs (**Figures 1E and 1F**). MLP protein levels were also lower in the proband EHTs, as expected^6^. These results revealed that calsarcin-1 is deficient in MLP-W4R;MYH7-R723C HCM proband heart and derived iPSC-CMs and EHTs.

**Figure 1.**
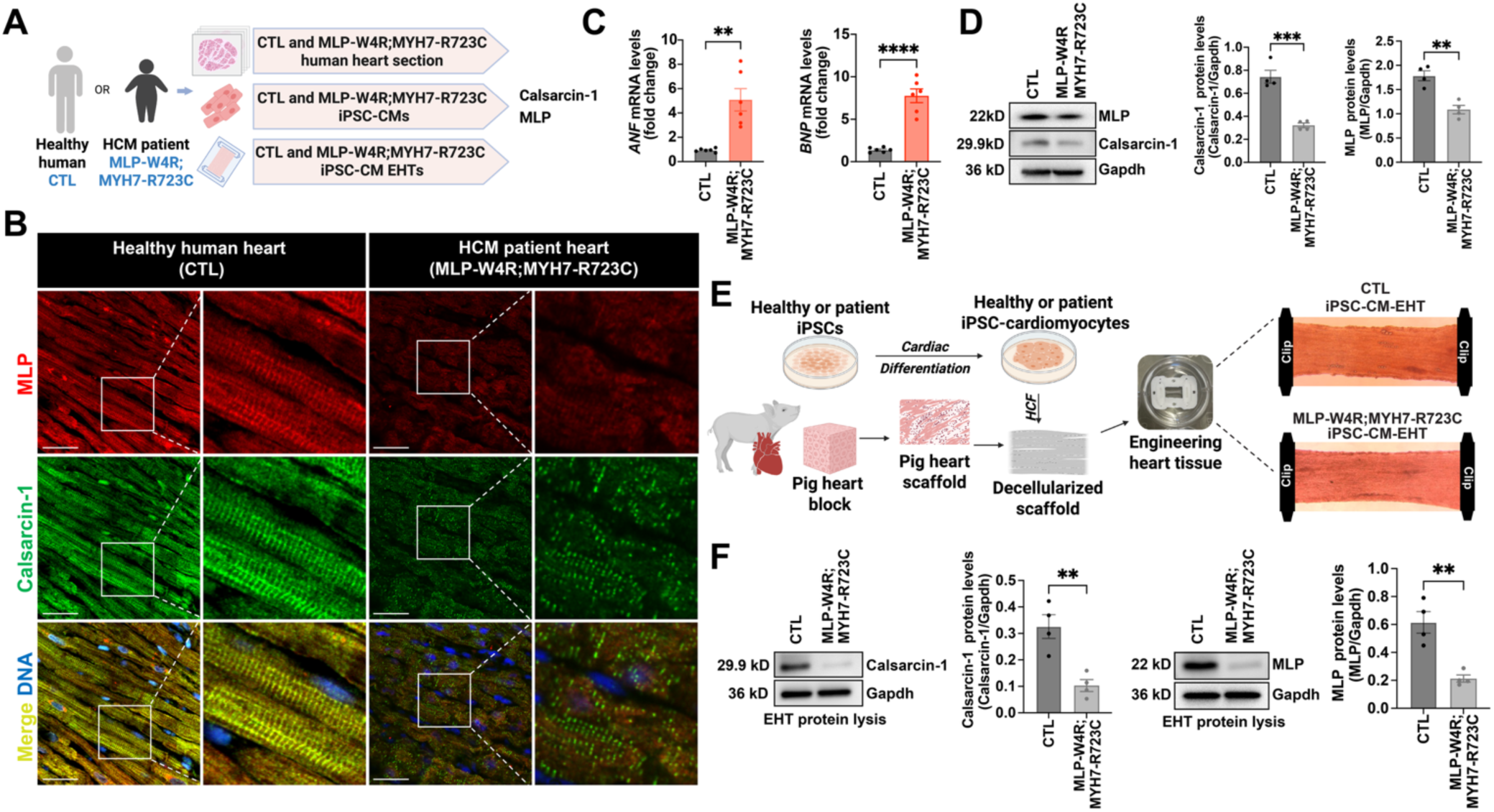
Expression of the muscle LIM protein (MLP) and calsarcin-1 in the healthy control (CTL) and proband cardiomyocytes. **A.** Schematic for the study. **B.** Representative immunofluorescence staining of calsarcin-1 and MLP in healthy human and MLP-W4R;MYH-R723C hypertrophic cardiomyopathy (HCM) patient heart sections (scale bar = 100 µm). **C.** Atrial natriuretic factor (*ANF*) and brain natriuretic peptide (*BNP*) mRNA levels in CTL and MLP-W4R;MYH-R723C iPSC-CMs, *n* = 6 biological replicates, with each replicate representing an independent batch of iPSC-CMs differentiation, \*\**P*<0.01, \*\*\*\**P*<0.0001 (2-tailed unpaired Student t test). **D.** Calsarcin-1 and MLP protein levels in CTL and MLP-W4R;MYH-R723C iPSC-CMs, *n*=4 independent batches of iPSC-CM differentiation, \*\**P*<0.01, \*\*\**P*<0.001 (2-tailed unpaired Student’s t-test). **E.** The flowchart for engineered heart tissue (EHT) generation and images of CTL and MLP-W4R;MYH-R723C iPSC-CM-EHTs. Each EHT was generated by seeding 1.35 × 10⁶ iPSC-CMs and 0.15 × 10⁶ human cardiac fibroblasts onto a decellularized porcine cardiac scaffold, followed by 7 days of static culture at a fixed length in a standard incubator without electrical pacing. **F.** Calsarcin-1 and MLP protein levels in CTL and MLP-W4R;MYH-R723C iPSC-CM-EHTs, *n*=4 independent EHT batches, each generated from an independent iPSC-CM differentiation, \*\**P*<0.01 (2-tailed unpaired Student’s t-test). All data are presented as mean ± SEM.

### Alleviation of hypertrophy by calsarcin-1 and its requirement in the mitigation of HCM defects by MLP in patient iPSC-derived cardiomyocytes

To investigate whether reduced calsarcin-1 in the MLP-W4R;MYH7-R723C HCM proband relates to MLP-dependent Z-disc mechanosensing (MZM) complex stability that modulates HCM, calsarcin-1 was ectopically expressed in CTL and MLP-W4R;MYH7-R723C iPSC-CMs via lentiviral transduction (**Figure 2A**). We found that calsarcin-1 overexpression significantly rescued the HCM phenotype in MLP-W4R;MYH7-R723C iPSC-CMs, including elevated *ANF* (**Figure 2B**) and *BNP* mRNA (**Figure 2C**) expression and enlarged cardiomyocyte sizes (**Figures 2D and 2E**), without affecting CTL iPSC-CMs. These findings suggest that ectopic calsarcin-1 could attenuate the hypertrophic phenotype of MLP-W4R:MYH7-R723C iPSC-CMs.

**Figure 2.**
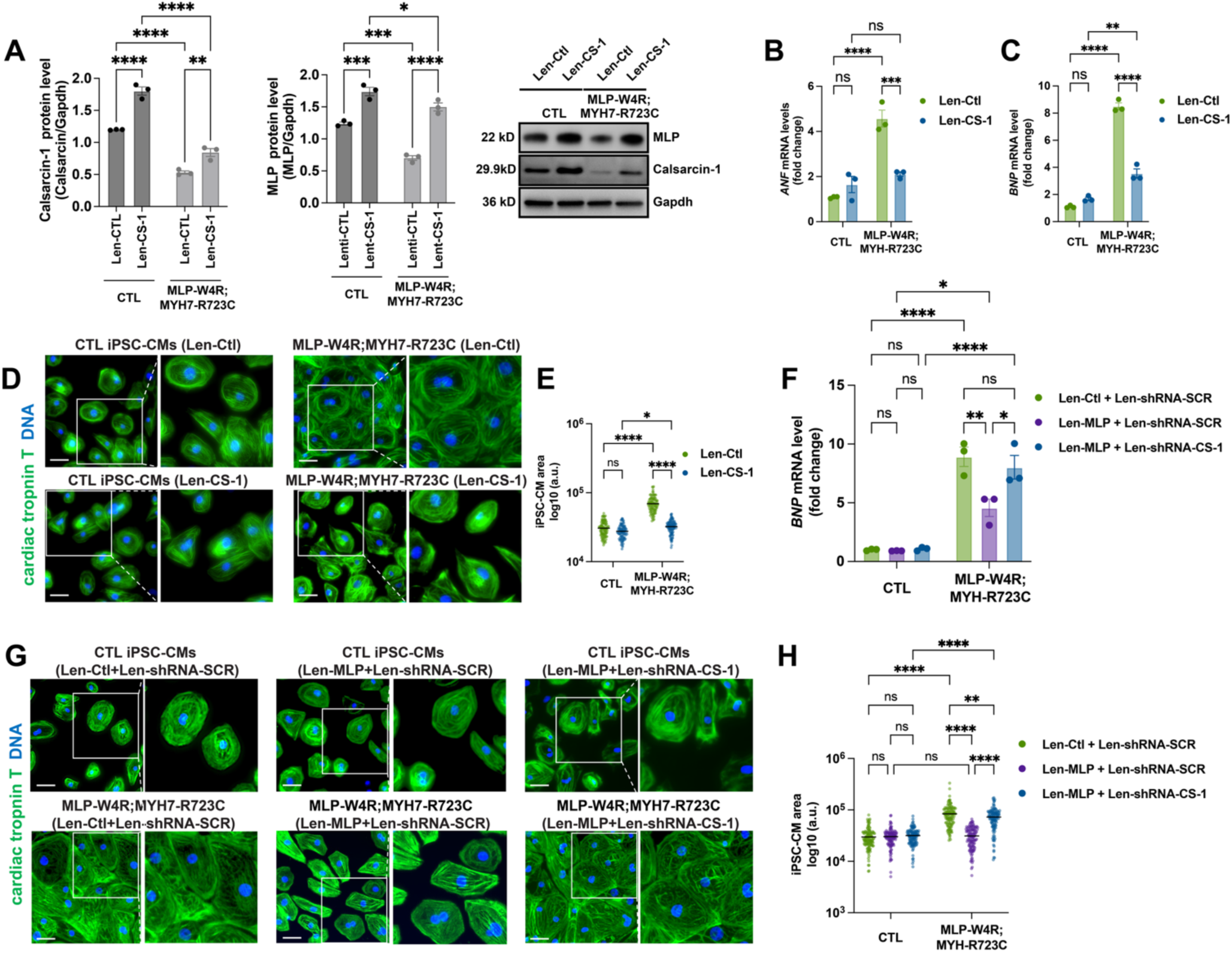
Regulation of hypertrophic defects by calsarcin-1 and MLP in patient-specific iPSC-derived cardiomyocytes. **A.** Calsarcin-1 and MLP protein levels in CTL and MLP-W4R;MYH-R723C iPSC-CMs treated with lentivirus expressing calsarcin-1 (Len-CS-1) or control lentivirus lacking calsarcin-1 expression (Len-CTL), *n*=3 independent batches of cardiac differentiation, \**P*<0.05, \*\**P*<0.01, \*\*\**P*<0.001, \*\*\*\**P*<0.0001, ns: not significant (Two-way ANOVA followed by Tukey multiple comparisons test; Calsarcin-1: interaction *P*= 0.0135; MLP: interaction *P*= 0.0178). **B-C**. Atrial natriuretic factor (*ANF)* and B-type natriuretic peptide (*BNP)* mRNA levels in CTL and MLP-W4R;MYH-R723C iPSC-CMs treated with Len-CS-1 or Len-Ctl, *n*=3 independent batches of cardiac differentiation, \*\**P*<0.01, \*\*\**P*<0.001, \*\*\*\**P*<0.0001, ns: not significant (Two-way ANOVA followed by Tukey multiple comparisons test; *ANF*: interaction *P* = 0.0005; *BNP*: interaction *P* < 0.0001). **D-E.** Cell sizes determined in CTL and MLP-W4R;MYH-R723C iPSC-CMs transduced with Len-CS-1 or Len-Ctl via cardiac troponin T (cTnT) and DAPI immunofluorescence staining (scale bar = 50 μm) and quantification of cell areas (in total 100-120 cells were calculated from three (*n*=3) independent batches of cardiac differentiation, ∼30-40 cells in each group per batch), \*\**P*<0.01, \*\*\*\**P*<0.0001, ns: not significant (Two-way ANOVA followed by Tukey multiple comparisons test; interaction *P* <0.0001). **F-H**. *BNP* mRNA level, representative cTnT and DAPI staining images (scale bar = 50 μm), and quantification of cell size in CTL and MLP-W4R;MYH-R723C iPSC-CMs infected with lentivirus expressing MLP (Len-MLP) or control lentivirus lacking MLP expression (Len-Ctl), lentivirus-shRNA-calsarcin-1 (Len-shRNA-CS-1), or lentivirus-shRNA-scramble (Len-shRNA-SCR), *n*=3 independent batches of cardiac differentiation, \*\**P*<0.01, \*\*\**P*<0.001, \*\*\*\**P*<0.0001, ns: not significant (Two-way ANOVA followed by Tukey multiple comparisons test; *BNP*: interaction *P* = 0.0115; Cell size: interaction *P <0.0001*). All data are presented as mean ± SEM.

We have previously reported a significant reduction in sarcomeric MLP levels in the heart of the MLP-W4R;MYH7-R723C proband, leading to disinhibition of calcineurin/NFAT signaling and consequent cardiac hypertrophy^6^. We therefore tested MLP expression levels in calsarcin-1 lentivirus-infected MLP-W4R;MYH7-R723C iPSC-CMs. Notably, lentiviral expression of calsarcin-1 significantly enhanced MLP expression in MLP-W4R;MYH7-R723C iPSC-CMs, compared with MLP-W4R;MYH7-R723C iPSC-CMs treated with Len-Ctl (**Figure 2A**). These results suggest that calsarcin-1 alleviated the hypertrophic defects of MLP-W4R;MYH7-R723C iPSC-CMs in part via stabilizing MLP within the MZM complex. To determine whether calsarcin-1 is required for MLP mechanosensing to rescue hypertrophic defects in MLP-W4R;MYH7-R723C iPSC-CMs, lentivirus-based shRNA targeting calsarcin-1 (Len-shRNA-CS-1) and lentivirus carrying a scramble sequence (Len-shRNA-SCR) were generated (**Figure S1**). We found that the effects of MLP in reducing both *BNP* mRNA (**Figure 2F**) and cell size (**Figure 2G**–**2H**) in MLP-W4R;MYH7-R723C iPSC-CMs were markedly blunted when calsarcin-1 was knocked down via Len-shRNA-CS-1, suggesting that calsarcin-1 is an essential regulator of MZM signaling in modulating HCM phenotypes.

### Attenuation of hypertrophic defects by calsarcin-1 in cardiomyocytes derived from additional HCM families

To determine whether an ectopic expression of calsarcin-1 could possibly alleviate hypertrophic cardiomyopathic defects caused by pathogenic mutations in sarcomeric genes other than MLP-W4R;MYH7-R723C in the proband, we investigated two additional iPSC lines generated from HCM patients carrying pathogenic variants in MYH7-R663H and MYBPC3-V321M (myosin-binding protein C)^13,14^. Calsarcin-1 was ectopically expressed in cardiomyocytes derived from these two HCM lines via lentiviral transduction. Notably, calsarcin-1 was able to significantly rescue the hypertrophic defects, including the enlarged cell size and abnormal *BNP* mRNA expression levels, in cardiomyocytes derived from the MYH7-R663H and MYBPC3-V321M (**Figure 3**) iPSC lines. These results suggest that calsarcin-1 may have a broader role in mitigating hypertrophic cardiomyopathic defects and could be an attractive therapeutic target for future HCM treatment.

**Figure 3.**
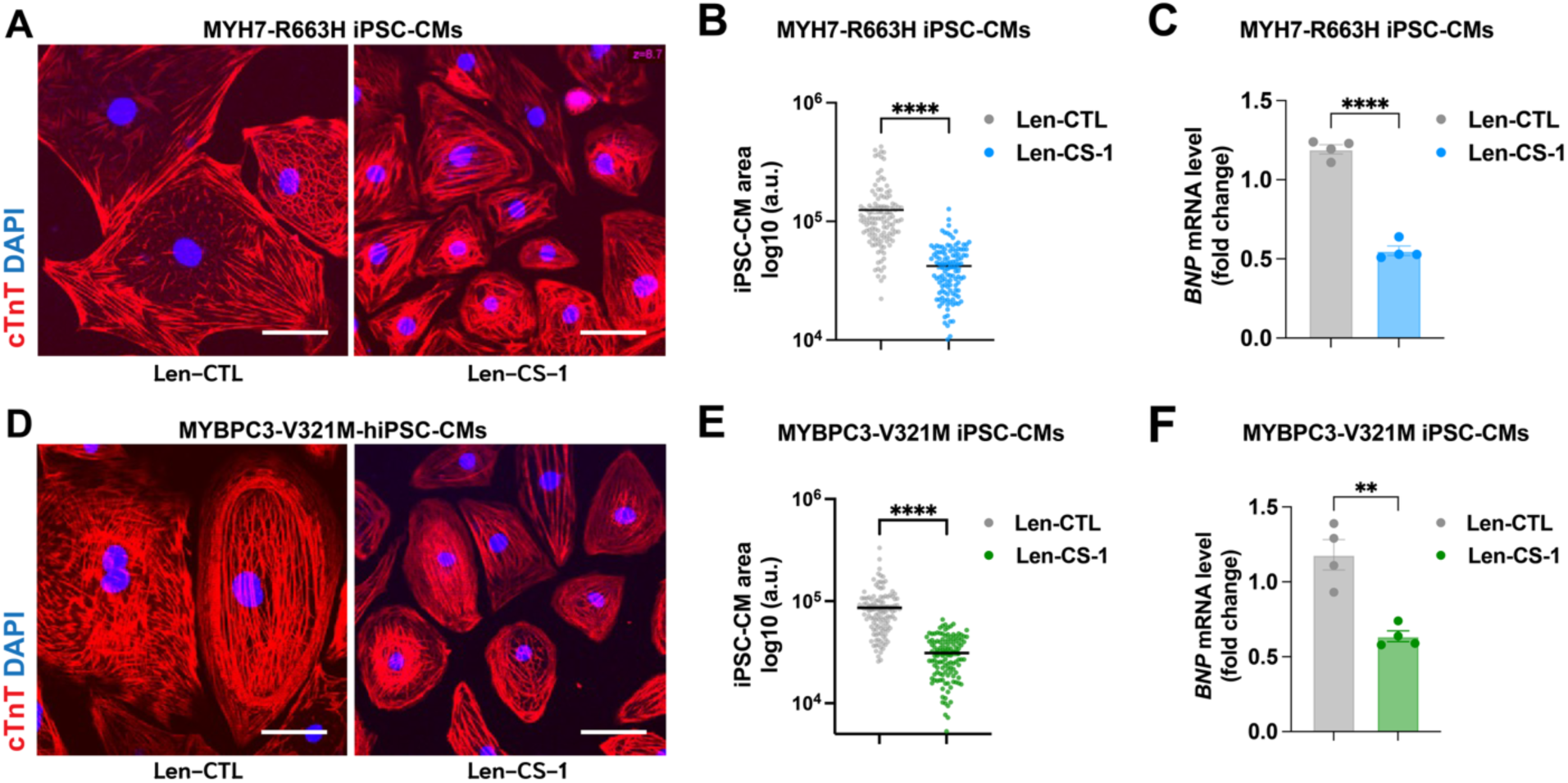
Ectopic expression of calsarcin-1 alleviated hypertrophy phenotypes in patient-specific iPSC-CMs carrying MYH7-R663H or MyBPC3-V321M. A-B. Representative images of cardiac troponin T (cTnT) immunofluorescence staining and cardiomyocyte size in MYH7-R663H iPSC-CMs infected with Len-CS-1 and Len-CTL, *n*=4 independent batches of cardiac differentiation, \*\*\*\**P*<0.0001 (2-tailed unpaired Mann-Whitney U test). **C.** The *BNP* mRNA expression level of MYH7-R663H iPSC-CMs with Len-CTL and Len-CS-1. *n*=4 independent batches of cardiac differentiation, \*\*\*\**P*<0.0001 (2-tailed unpaired Student’s t*-*test) **D-E.** Representative images based on cTnT immunofluorescence staining and CM area as a measure of cell size were presented for MyBPC3-V321M iPSC-CMs. The MyBPC3-V321M iPSC-CMs were infected with lentiviruses expressing either Len-CTL or Len-CS-1, *n*=4 independent batches of cardiac differentiation, \*\*\*\**P*<0.0001 (2-tailed unpaired Mann-Whitney U test). F. *BNP* mRNA expression levels in MyBPC3-V321M iPSC-CMs with Len-CTL and Len-CS-1 *n*=4 independent batches of cardiac differentiation, *n*=4 independent batches of cardiac differentiation, \*\**P*<0.01 (2-tailed unpaired Student’s t-test). All data are presented as mean ± SEM; ns: not significant. *MYH7*: myosin heavy chain 7; *MyBPC3*: myosin binding protein C.

### Calsarcin-1 functions as an important regulator of MLP/Z-disc mechanosensing in engineered heart tissues trained under different loading conditions

To further investigate the role of calsarcin-1 in MLP/Z-disc mechanosensing under disease-relevant mechanical stress, we developed a novel dynamic/isometric electrical-pacing bioreactor that enables chronic exposure of iPSC-CM-EHTs to precisely controlled loading conditions (**Figures 4A and 4B**). For this study, the bioreactor was programmed to apply two distinct types of synchronized mechanical and electrical stimulation to EHTs: A low-afterload condition consisting of cyclic 10% shortening at 1 Hz (**Figures 4B and 4C**), and a high-afterload condition consisting of maintaining EHTs under static tension without mechanical displacement while applying 1 Hz electrical pacing (**Figures 4C and 4D**). These regimens enabled reproducible modeling of physiological and pressure-overloaded mechanical environments, respectively.

**Figure 4.**
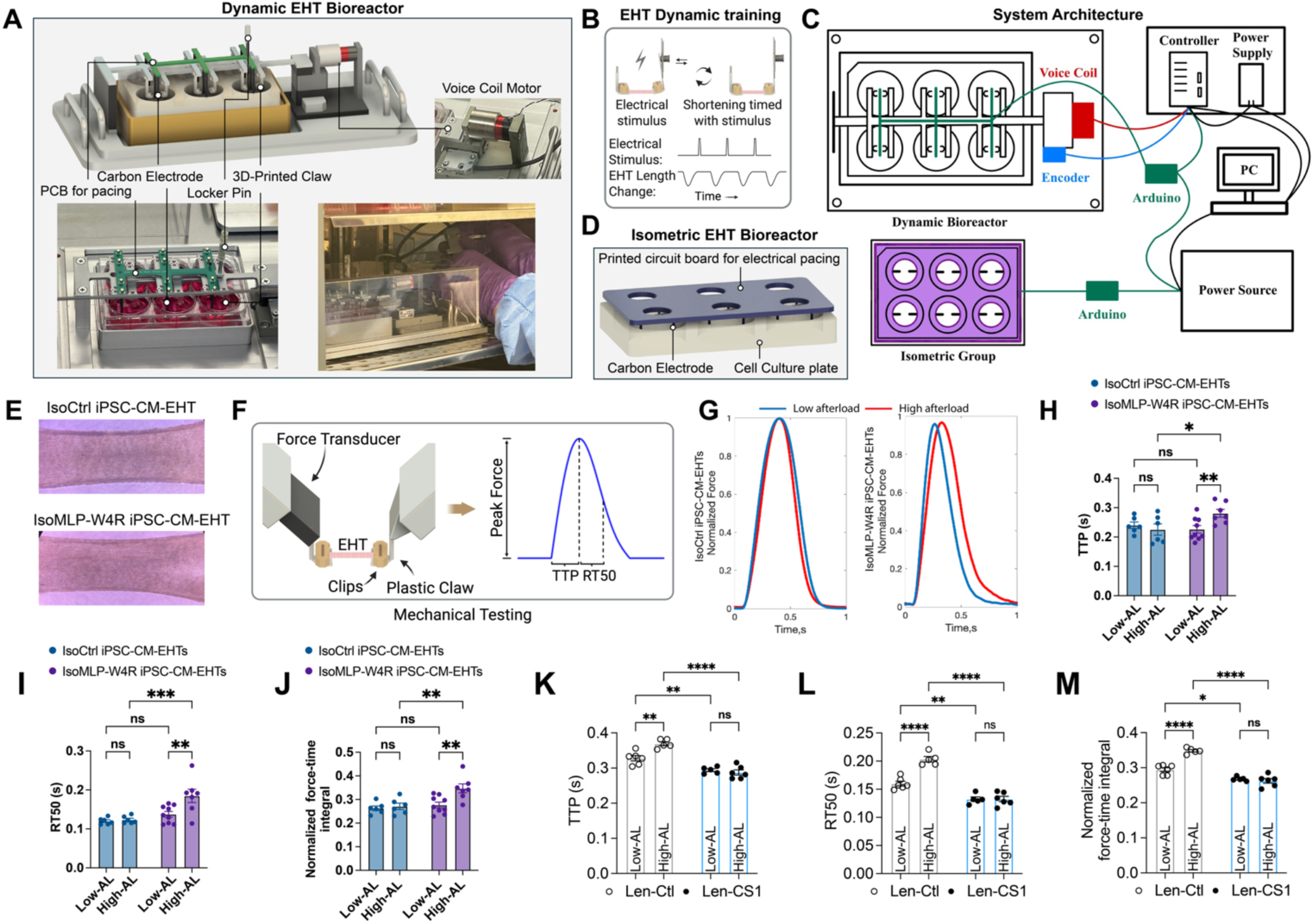
Ectopic expression of calsarcin-1 rescued dysfunction in IsoMLP-W4R iPSC-cardiomyocyte-engineered heart tissues exposed to pathological high afterload. **A**. The dynamic bioreactor applies cyclic uniaxial strain to engineered heart tissues (EHTs) using a voice coil actuator guided by a precision rail. Motion is tracked by an LM10 magnetic encoder and controlled via a Galil DMC-4103 system executing synchronized motion and pacing profiles. The system is housed in a sterile, incubator-compatible chamber. **B-C**. Schematic overview of the integrated dynamic/isometric EHTs bioreactor culture system. The platform comprises a dynamic bioreactor and an isometric culture module controlled by an Arduino-based controller that coordinates a voice-coil actuator and an encoder through a closed-loop feedback system. A dedicated power supply drives the actuation and control electronics, while a PC interface enables system programming, real-time movement monitoring, and electrical pacing, thereby providing precise mechanical conditioning of engineered heart tissues under tunable loading conditions. **D**. In high afterload EHT culture, the tissue is typically maintained in an isometric state where it generates force against a fixed length with minimal shortening in a 6-well plate covered with a pacing board equipped with carbon electrodes for electrical stimulation. **E.** Representative image of IsoCtrl iPSC-CM-EHTs and IsoMLP-W4R iPSC-CM-EHTs generated from isogenic control iPSC (IsoCtrl-iPSC, both MLP-W4R and MYH-R723C gene-corrected) and IsoMLP-W4R iPSC (MYH-R723C single correction) derived CMs. **F**. EHT functional assessment platform. Contractile and relaxation kinetics were evaluated using a high-sensitivity force transducer to determine peak force, time to peak force (TTP), and time from peak force to 50% relaxation (RT50), providing integrated measures of tissue systolic and diastolic performance. **G-J**. Average Force-Time curve, TTP, RT50, and normalized force-time integral (nFTI) of IsoCtrl iPSC-CM-EHTs and IsoMLP-W4R iPSC-CM-EHTs measured after 7 days of culturing in dynamic/isometric EHT bioreactors. Six independent EHT batches included both IsoCtrl and IsoMLP-W4R iPSC-CM-EHTs under low- and high-afterload conditions, with three additional independent batches of IsoMLP-W4R iPSC-CM-EHTs analyzed under both conditions (*n* = 6 for IsoCtrl; *n* = 9 for IsoMLP-W4R). Each batch was generated from an independent iPSC-CM differentiation; \**P*<0.05, \*\**P*<0.01, \*\*\**P*<0.001, ns: not significant (Two-way ANOVA followed by Tukey multiple comparisons test; TTP: interaction *P* = 0.0259, RT50: interaction *P* = 0.0427, and nFTI: interaction *P* = 0.0430). **K-M.** TTP, RT50, and nFTI were assessed in IsoMLP-W4R iPSC-CM-EHTs following lentiviral ectopic expression of calsarcin-1 (Len-CS-1) or a lentivirus lacking calsarcin-1 as a control (Len-Ctl), *n*=6 independent experimental batches, \**P*<0.05, \*\**P*<0.01, \*\*\*\**P*<0.0001, ns: not significant (Two-way ANOVA followed by Tukey multiple comparisons test; TTP: interaction *P* = 0.0020, RT50: interaction *P* = 0.0001, and nFTI: interaction *P* <0.0001). All data are presented as mean ± SEM.

These high- and low-afterload conditions were administered to EHTs with WT MLP (IsoCtrl-iPSCs, isogenic control iPSCs made through gene correction of both MLP and MYH7 mutations in the MLP-W4R;MYH7-R723C proband iPSCs) and mutant MLP (IsoMLP-W4R-iPSCs, isogenic MLP-W4R iPSCs made by correcting only the MYH7 mutation in the proband’s iPSCs; **Figures S2 A-D**). IsoCtrl iPSCs and IsoMLP-W4R iPSCs with normal chromosomal integrity and pluripotency were selected and expanded for cardiac differentiation (**Figures S2 E-F**). We further compared cardiomyocytes derived from the isogenic control (IsoCtrl iPSC-CMs) with those derived from an independent healthy donor iPSC line (CTL iPSC-CMs), and observed similar expression of calsarcin-1 and MLP proteins, *ANF* and *BNP* mRNA levels, and cardiomyocyte sizes, suggesting a normal phenotype of IsoCtrl iPSC-CMs (**Figure S3**).

The IsoCtrl iPSC-CM-EHTs and IsoMLP-W4R iPSC-CM-EHTs were generated and cultured in the dynamic/isometric EHT bioreactor for 7 days (**Figure 4E**). After dynamic culture, EHTs were removed, and their systolic and diastolic functions were evaluated (**Figures 4F-4G**). The additional mechanical resistance imposed by high afterload resulted in a delay in the time required to reach peak force (time to peak force, or TTP, **Figure 4H**). Slowed relaxation kinetics, seen as an increase in the time from peak force to 50% relaxation (RT50), were observed in IsoMLP-W4R iPSC-CM-EHTs under high afterload compared with the same EHTs under low afterload conditions and compared with IsoCtrl iPSC-CM-EHTs (**Figure 4I**). The normalized force-time integral (nFTI), representing the cumulative force generated over a single beat, was also increased in IsoMLP-W4R iPSC-CM-EHTs. In contrast, there was no significant change in the nFTI of high afterload IsoCtrl iPSC-CM-EHTs (**Figure 4J**).

These experiments were repeated with ectopic calsarcin-1 expression via lentiviral transduction (Len-CS1) in IsoMLP-W4R iPSC-CM-EHTs to assess whether the effects of high afterload were potentially mediated by insufficient calsarcin-1 function. Len-CS1 treatment of MLP-W4R iPSC-CM-EHTs prevented significant increases in TTP, RT50, and nFTI in response to high afterload, whereas those treated with a lentivirus lacking calsarcin-1 expression (Len-Ctl) exhibited the same systolic and diastolic defects as before (**Figures 4K-4M**). These findings are consistent with the role of calsarcin-1 as a mechanistically important regulator of pathological cardiac remodeling, potentially by enhancing the MLP/Z-disc complex, and highlight calsarcin-1 as a potential therapeutic target in hypertrophic cardiomyopathy.

## DISCUSSION

Hypertrophic cardiomyopathy is the most common cardiac disorder and a leading cause of sudden cardiac death in the young^15^. Pathogenic variants in MYH7 and MYBPC3 are the common genetic causes among genotype-positive cases; however, the mechanistic link among primary sarcomere defects, downstream molecular remodeling, and the ultimate hypertrophic phenotype remains incompletely understood^16,17^. We hypothesized that a co-segregating CSRP3 (encoding muscle LIM protein, MLP) missense (W4R) in a proband harboring the MYH7-R723C mutation would reveal a previously unrecognized mechanistic node linking sarcomere dysfunction to hypertrophic remodeling.

Herein, we found that calsarcin-1 expression was consistently reduced in the MLP-W4R;MYH-R723C proband myocardium, patient-specific iPSC-CMs, and EHTs, and that it closely correlated with Z-disc disorganization, suggesting that loss of calsarcin-1 is an important molecular signature of sarcomere destabilization. Notably, restoring calsarcin-1 via lentiviral delivery enhanced MLP expression, reversed cardiomyocyte hypertrophy, and normalized *ANF* and *BNP* expression, whereas calsarcin-1 depletion by shRNA abrogated the anti-hypertrophic benefit of ectopic MLP expression in proband CMs. Importantly, calsarcin-1 gain-of-function also rescued hypertrophic defects in CMs from two additional HCM families (MYH7-R663H and MYBPC3-V321M), suggesting it may have a broad role in mitigating pathological hypertrophy and serve as a key effector linking MLP/Z-disc biomechanical sensing to modulation of hypertrophic signaling. Moreover, using isogenic control and MLP-W4R iPSC-derived CMs and EHTs as well as a newly developed EHT bioreactor platform that integrates controlled afterload with synchronized electrical pacing, we showed that ectopic calsarcin-1 expression restored sarcomeric regulation in IsoMLP-W4R iPSC-CM-EHTs even under pathological mechanical stress, insofar as it prevented changes in TTP, RT50, and nFTI. Together, these findings define calsarcin-1 as a previously unrecognized mechanosensitive modifier that couples MLP/Z-disc integrity to cellular hypertrophy and contractile misregulation, and establish an important therapeutic target whereby restoration of calsarcin-1, a critical Z-disc component, can reverse MLP-W4R-driven hypertrophic remodeling under high afterload.

Among Z-disc-resident LIM proteins, MLP is essential for cytoarchitectural integrity. Heterozygous and biallelic CSRP3 variants contribute to hypertrophic cardiomyopathy^18,19^. The W4R variant localizes to the N-terminal LIM1 domain required for direct α-actinin-2 binding; impaired interaction compromises MLP Z-disc anchoring, accelerates proteasomal turnover, and renders cardiomyocytes vulnerable to biomechanical stress^20–22^. As previously demonstrated, MLP-W4R likely acts as a partial loss-of-function allele that, in combination with MYH7-R723C, establishes a heightened mechanobiological stress state characterized by accelerated MLP depletion and impaired downstream signaling^6^. The marked loss of MLP in proband myocardium, iPSC-CMs, and EHTs validates this model and offers insights into the severe clinical phenotype.

Calsarcin-1 is the predominant cardiac isoform of the Calsarcin family and serves as a critical Z-disc organizer by binding α-actinin and calcineurin into a functional macromolecular complex^7,8,23,24^. Loss of calsarcin-1 destabilizes the Z-disc scaffold, promoting maladaptive hypertrophic signaling through both calcineurin–NFAT and Hippo–YAP/TAZ pathways^8,25–28^. Our findings expand the current model by identifying calsarcin-1 insufficiency as a convergent defect in MLP-W4R;MYH-R723C HCM pathogenesis and reinforce the concept that Z-disc complex integrity, rather than MLP expression alone, determines MLP-dependent signaling output. As gain-of-function of calsarcin-1 via lentiviral expression elevated MLP expression and mitigated hypertrophy in proband CMs, it is likely that MLP and calsarcin-1 may enhance each other at the Z-disc, enabling MLP-mediated mechano-protection in cardiomyocytes.

Monolayer iPSC-CMs recapitulate many HCM features^13^, but three-dimensional EHTs are required to interrogate sarcomere-level contractile phenotypes under defined biomechanical conditions^14,29^. Our EHT training bioreactors add independent modulation of mechanical afterload and synchronous electrical pacing, enabling defined physiological and pathological afterload regimens to be delivered to the EHTs over time. Under high afterload, MLP-W4R iPSC-CM-EHTs developed a hypercontractile phenotype characterized by prolonged TTP, RT50, and increased nFTI, a signature consistent with hyperdynamic diastolic dysfunction in HCM patient^15,16^. The differential benefit of ectopic expression of calsarcin-1 under high versus low afterload reveals a load-dependent role for the calsarcin-1/Z-disc complex. Rather than serving as a sole structural component, calsarcin-1 emerges as a mechanosensing component that becomes functionally limiting only under pathological mechanical stress, consistent with the impaired cardiac adaptation to increased hemodynamic stress and cardiomyopathic remodeling observed in MLP-deficient mice^4,30,31^. Our data advance the concept of the Z-disc beyond a structural anchor, confirming the notion that it is also a dynamic mechanosensory hub that determines cardiomyocyte resilience and vulnerability under biomechanical stress^32^.

These findings identify calsarcin-1 as a potential therapeutic target for HCM. Although cardiac myosin inhibitors, including mavacamten and aficamten, have shown benefit in modulating sarcomere mechanics in obstructive HCM^33^, they do not directly address the upstream Z-disc mechanosensing pathways that drive maladaptive hypertrophic remodeling. Restoring Z-disc signaling integrity through calsarcin-1 enhancement may represent a complementary disease-modifying strategy. As AAV-based targeting of Z-disc proteins has shown promise in preclinical dilated cardiomyopathy models^34^, it would be of significant interest to extend this approach to assess whether ectopic expression of calsarcin-1 can mitigate HCM based on our findings herein.

## Limitations of the study

Some limitations should be noted. Although our data defines a critical role for calsarcin-1 in the MLP/Z-disc complex, future endeavors are warranted to dissect downstream signaling pathways responsible for its effects. Future efforts will also be made to perform transcriptomic and phospho-proteomic studies to identify key mechanotransduction effectors, including potential involvement of calcineurin-NFAT, Hippo-YAP/TAZ, and MAPK pathways. Moreover, while calsarcin-1 is required for MLP-mediated suppression of hypertrophic remodeling, our studies do not exclude its additional MLP-independent functions. Investigation of calsarcin-1 functions in MLP deficient models may help provide further insights into distinct contributions of these mechanisms in the future.

## CONCLUSION

This study identifies calsarcin-1 as a previously unrecognized molecular determinant of MLP/Z-disc complex integrity in HCM. Therapeutic restoration of calsarcin-1 enhances MLP/Z-disc stability and attenuates pathological cardiac remodeling. Furthermore, patient-specific EHTs subjected to pathological HCM loading in a bioreactor provide functional validation of calsarcin-1 as a promising therapeutic target for HCM.

## RESOURCE AVAILABILITY

### Lead contact

Further information and requests should be directed to and will be fulfilled by the lead contact, Professor Yibing Qyang.

## Materials availability

All iPSC lines generated in this study are available from the Lead Contact upon reasonable request.

## Data and code availability

- The data reported in this paper will be shared by the lead contact upon reasonable request.
- This paper does not report original code.
- Any additional information required to reanalyze the data reported in this work is available from the lead contact upon request.

## ACKNOWLEDGMENTS

This work was supported by the National Institutes of Health (NIH) R01HL171984 and R01HL164783 to Dr. Yibing Qyang, and NIH R01HL163092 to Dr. Stuart G. Campbell.

## AUTHOR CONTRIBUTIONS

Xin Li, S.G. Campbell, and Y. Qyang conceived the study; Xin Li, Y. Lu and Xianmu Li performed research; Xin Li and Y. Qyang wrote the manuscript. Xin Li, Y. Lu and Xianmu Li, M. Riaz, M. Hoareau, S. G. Campbell and Y. Qyang edited the manuscript. Thomas Blum guided and advised on bioreactor protocol; M.Riaz and J. Park guided and advised on iPSC-derived cardiomyocyte differentiation and gene editing. H.Q. Luo, and Y. Lu assisted in iPSC-derived cardiomyocyte differentiation M. Hoareau assisted in cardiomyocyte identity verification. Ilhan Gokhan, Xia Li guided and advised on EHT seeding, culturing and functional testing; S. Fooladi and Q. Xing contributed to scientific discussions regarding the project; Y. Qyang, S.G. Campbell, and L. Yue provided valuable input and guidance during manuscript preparation and submission.

## DECLARATION OF INTERESTS

Dr. Stuart G. Campbell is co-founder of and has an ownership stake in Propria Inc., which has licensed EHT technology used in this study. The other authors declare no conflicts of interest.

## References

1. Maron BJ, Estes NA, 3rd, Maron MS, Almquist AK, Link MS, Udelson JE. Primary prevention of sudden death as a novel treatment strategy in hypertrophic cardiomyopathy. Circulation. 2003;107:2872–2875. doi: 10.1161/01.CIR.0000072343.81530.75

2. Seidman JG, Seidman C. The genetic basis for cardiomyopathy: from mutation identification to mechanistic paradigms. Cell. 2001;104:557–567. doi: 10.1016/s0092-8674(01)00242-2

3. Green EM, Wakimoto H, Anderson RL, Evanchik MJ, Gorham JM, Harrison BC, Henze M, Kawas R, Oslob JD, Rodriguez HM, et al. A small-molecule inhibitor of sarcomere contractility suppresses hypertrophic cardiomyopathy in mice. Science. 2016;351:617–621. doi: 10.1126/science.aad3456

4. Arber S, Hunter JJ, Ross J, Jr., Hongo M, Sansig G, Borg J, Perriard JC, Chien KR, Caroni P. MLP-deficient mice exhibit a disruption of cardiac cytoarchitectural organization, dilated cardiomyopathy, and heart failure. Cell. 1997;88:393–403. doi: 10.1016/s0092-8674(00)81878-4

5. Knoll R, Hoshijima M, Hoffman HM, Person V, Lorenzen-Schmidt I, Bang ML, Hayashi T, Shiga N, Yasukawa H, Schaper W, et al. The cardiac mechanical stretch sensor machinery involves a Z disc complex that is defective in a subset of human dilated cardiomyopathy. Cell. 2002;111:943–955. doi: 10.1016/s0092-8674(02)01226-6

6. Riaz M, Park J, Sewanan LR, Ren Y, Schwan J, Das SK, Pomianowski PT, Huang Y, Ellis MW, Luo J, et al. Muscle LIM Protein Force-Sensing Mediates Sarcomeric Biomechanical Signaling in Human Familial Hypertrophic Cardiomyopathy. Circulation. 2022;145:1238–1253. doi: 10.1161/CIRCULATIONAHA.121.056265

7. Frey N, Richardson JA, Olson EN. Calsarcins, a novel family of sarcomeric calcineurin-binding proteins. Proc Natl Acad Sci U S A. 2000;97:14632–14637. doi: 10.1073/pnas.260501097

8. Frey N, Barrientos T, Shelton JM, Frank D, Rutten H, Gehring D, Kuhn C, Lutz M, Rothermel B, Bassel-Duby R, et al. Mice lacking calsarcin-1 are sensitized to calcineurin signaling and show accelerated cardiomyopathy in response to pathological biomechanical stress. Nat Med. 2004;10:1336–1343. doi: 10.1038/nm1132

9. Vikstrom KL, Bohlmeyer T, Factor SM, Leinwand LA. Hypertrophy, pathology, and molecular markers of cardiac pathogenesis. Circ Res. 1998;82:773–778. doi: 10.1161/01.res.82.7.773

10. Hasegawa K, Fujiwara H, Doyama K, Miyamae M, Fujiwara T, Suga S, Mukoyama M, Nakao K, Imura H, Sasayama S. Ventricular expression of brain natriuretic peptide in hypertrophic cardiomyopathy. Circulation. 1993;88:372–380. doi: 10.1161/01.cir.88.2.372

11. Geske JB, McKie PM, Ommen SR, Sorajja P. B-type natriuretic peptide and survival in hypertrophic cardiomyopathy. J Am Coll Cardiol. 2013;61:2456–2460. doi: 10.1016/j.jacc.2013.04.004

12. Schwan J, Kwaczala AT, Ryan TJ, Bartulos O, Ren Y, Sewanan LR, Morris AH, Jacoby DL, Qyang Y, Campbell SG. Anisotropic engineered heart tissue made from laser-cut decellularized myocardium. Sci Rep. 2016;6:32068. doi: 10.1038/srep32068

13. Lan F, Lee AS, Liang P, Sanchez-Freire V, Nguyen PK, Wang L, Han L, Yen M, Wang Y, Sun N, et al. Abnormal calcium handling properties underlie familial hypertrophic cardiomyopathy pathology in patient-specific induced pluripotent stem cells. Cell stem cell. 2013;12:101–113. doi: 10.1016/j.stem.2012.10.010

14. Seeger T, Shrestha R, Lam CK, Chen C, McKeithan WL, Lau E, Wnorowski A, McMullen G, Greenhaw M, Lee J, et al. A Premature Termination Codon Mutation in MYBPC3 Causes Hypertrophic Cardiomyopathy via Chronic Activation of Nonsense-Mediated Decay. Circulation. 2019;139:799–811. doi: 10.1161/CIRCULATIONAHA.118.034624

15. Maron BJ, Maron MS. The 25-year genetic era in hypertrophic cardiomyopathy: revisited. Circulation Cardiovascular genetics. 2014;7:401–404. doi: 10.1161/CIRCGENETICS.114.000741

16. Marian AJ, Braunwald E. Hypertrophic Cardiomyopathy: Genetics, Pathogenesis, Clinical Manifestations, Diagnosis, and Therapy. Circ Res. 2017;121:749-770. doi: 10.1161/CIRCRESAHA.117.311059

17. Ho CY, Day SM, Ashley EA, Michels M, Pereira AC, Jacoby D, Cirino AL, Fox JC, Lakdawala NK, Ware JS, et al. Genotype and Lifetime Burden of Disease in Hypertrophic Cardiomyopathy: Insights from the Sarcomeric Human Cardiomyopathy Registry (SHaRe). Circulation. 2018;138:1387–1398. doi: 10.1161/CIRCULATIONAHA.117.033200

18. Geier C, Gehmlich K, Ehler E, Hassfeld S, Perrot A, Hayess K, Cardim N, Wenzel K, Erdmann B, Krackhardt F, et al. Beyond the sarcomere: CSRP3 mutations cause hypertrophic cardiomyopathy. Hum Mol Genet. 2008;17:2753–2765. doi: 10.1093/hmg/ddn160

19. Newman B, Cescon D, Woo A, Rakowski H, Erikkson MJ, Sole M, Wigle ED, Siminovitch KA. W4R variant in CSRP3 encoding muscle LIM protein in a patient with hypertrophic cardiomyopathy. Mol Genet Metab. 2005;84:374–375. doi: 10.1016/j.ymgme.2004.11.013

20. Lindholm ME, Jimenez-Morales D, Zhu H, Seo K, Amar D, Zhao C, Raja A, Madhvani R, Abramowitz S, Espenel C, et al. Mono- and Biallelic Protein-Truncating Variants in Alpha-Actinin 2 Cause Cardiomyopathy Through Distinct Mechanisms. Circulation Genomic and precision medicine. 2021;14:e003419. doi: 10.1161/CIRCGEN.121.003419

21. Klaavuniemi T, Alho N, Hotulainen P, Kelloniemi A, Havukainen H, Permi P, Mattila S, Ylanne J. Characterization of the interaction between Actinin-Associated LIM Protein (ALP) and the rod domain of alpha-actinin. BMC Cell Biol. 2009;10:22. doi: 10.1186/1471-2121-10-22

22. Atang AE, Rebbeck RT, Thomas DD, Avery AW. Cardiomyopathy-associated variants alter the structure and function of the alpha-actinin-2 actin-binding domain. Biochem Biophys Res Commun. 2023;670:12–18. doi: 10.1016/j.bbrc.2023.05.050

23. Frank D, Frey N. Cardiac Z-disc signaling network. J Biol Chem. 2011;286:9897–9904. doi: 10.1074/jbc.R110.174268

24. Pyle WG, Solaro RJ. At the crossroads of myocardial signaling: the role of Z-discs in intracellular signaling and cardiac function. Circ Res. 2004;94:296–305. doi: 10.1161/01.RES.0000116143.74830.A9

25. Paulsson AK, Franklin S, Mitchell-Jordan SA, Ren S, Wang Y, Vondriska TM. Post-translational regulation of calsarcin-1 during pressure overload-induced cardiac hypertrophy. J Mol Cell Cardiol. 2010;48:1206–1214. doi: 10.1016/j.yjmcc.2010.02.009

26. Knoll R, Linke WA, Zou P, Miocic S, Kostin S, Buyandelger B, Ku CH, Neef S, Bug M, Schafer K, et al. Telethonin deficiency is associated with maladaptation to biomechanical stress in the mammalian heart. Circ Res. 2011;109:758–769. doi: 10.1161/CIRCRESAHA.111.245787

27. Knoll R, Buyandelger B, Lab M. The sarcomeric Z-disc and Z-discopathies. Journal of biomedicine & biotechnology. 2011;2011:569628. doi: 10.1155/2011/569628

28. Purevjav E, Towbin JA. Cardiomyopathy - Disease of the Heart Muscle. 2021.

29. Carter RD, Purnama U, Castro-Guarda M, Montes-Aparicio CN, Chandran A, Mbasu R, Ruby M, Daly C, Brisk K, Christian HC, et al. Systems Biology and Functional Assessments of Human iPSC-Cardiomyocyte Models of Insulin Resistance Capture Key Hallmarks of Diabetic Cardiomyopathy. Diabetes. 2025;74:1929–1945. doi: 10.2337/db25-0204

30. Li X, Lu WJ, Li Y, Wu F, Bai R, Ma S, Dong T, Zhang H, Lee AS, Wang Y, et al. MLP-deficient human pluripotent stem cell derived cardiomyocytes develop hypertrophic cardiomyopathy and heart failure phenotypes due to abnormal calcium handling. Cell death & disease. 2019;10:610. doi: 10.1038/s41419-019-1826-4

31. Buyandelger B, Ng KE, Miocic S, Piotrowska I, Gunkel S, Ku CH, Knoll R. MLP (muscle LIM protein) as a stress sensor in the heart. Pflugers Arch. 2011;462:135–142. doi: 10.1007/s00424-011-0961-2

32. Lyon RC, Zanella F, Omens JH, Sheikh F. Mechanotransduction in cardiac hypertrophy and failure. Circ Res. 2015;116:1462–1476. doi: 10.1161/CIRCRESAHA.116.304937

33. Braunwald E, Saberi S, Abraham TP, Elliott PM, Olivotto I. Mavacamten: a first-in-class myosin inhibitor for obstructive hypertrophic cardiomyopathy. Eur Heart J. 2023;44:4622–4633. doi: 10.1093/eurheartj/ehad637

34. Balakrishnan B, Altassan R, Budhraja R, Liou W, Lupo A, Bryant S, Mankouski A, Radenkovic S, Preston GJ, Pandey A, et al. AAV-based gene therapy prevents and halts the progression of dilated cardiomyopathy in a mouse model of phosphoglucomutase 1 deficiency (PGM1-CDG). Translational research : the journal of laboratory and clinical medicine. 2023;257:1–14. doi: 10.1016/j.trsl.2023.01.004

35. Ho CY, Charron P, Richard P, Girolami F, Van Spaendonck-Zwarts KY, Pinto Y. Genetic advances in sarcomeric cardiomyopathies: state of the art. Cardiovasc Res. 2015;105:397–408. doi: 10.1093/cvr/cvv025

