## Supplemental information for "Calsarcin-1 modulates the muscle LIM protein/Z-disc complex mechanosensing in familial hypertrophic cardiomyopathy"

Document S1. Figures S1–S3

**Figure S1**

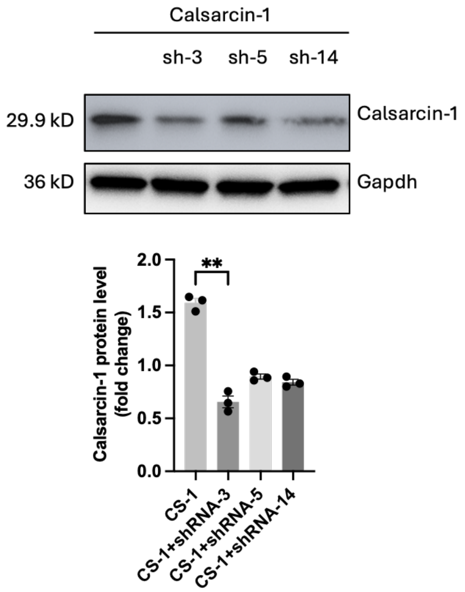


**Figure S1. Related to Figure 2. Evaluation of knockdown efficiency of calsarcin-1 shRNAs.** Three short hairpin RNAs (shRNAs) of calsarcin-1 were designed, including shRNA #3 (sh-3), #5 (sh-5), and #14 (sh-14). HEK293 cells overexpressing calsarcin-1 and co-transfected with shRNAs were used to assess calsarcin-1 protein expression levels by Western blot. (*n*=3 independent batches of infection and Western blot experiments). All data are presented as mean ± SEM; ***P*<0.01 (one-way ANOVA followed by Tukey's multiple-comparison tests).

**Figure S2**

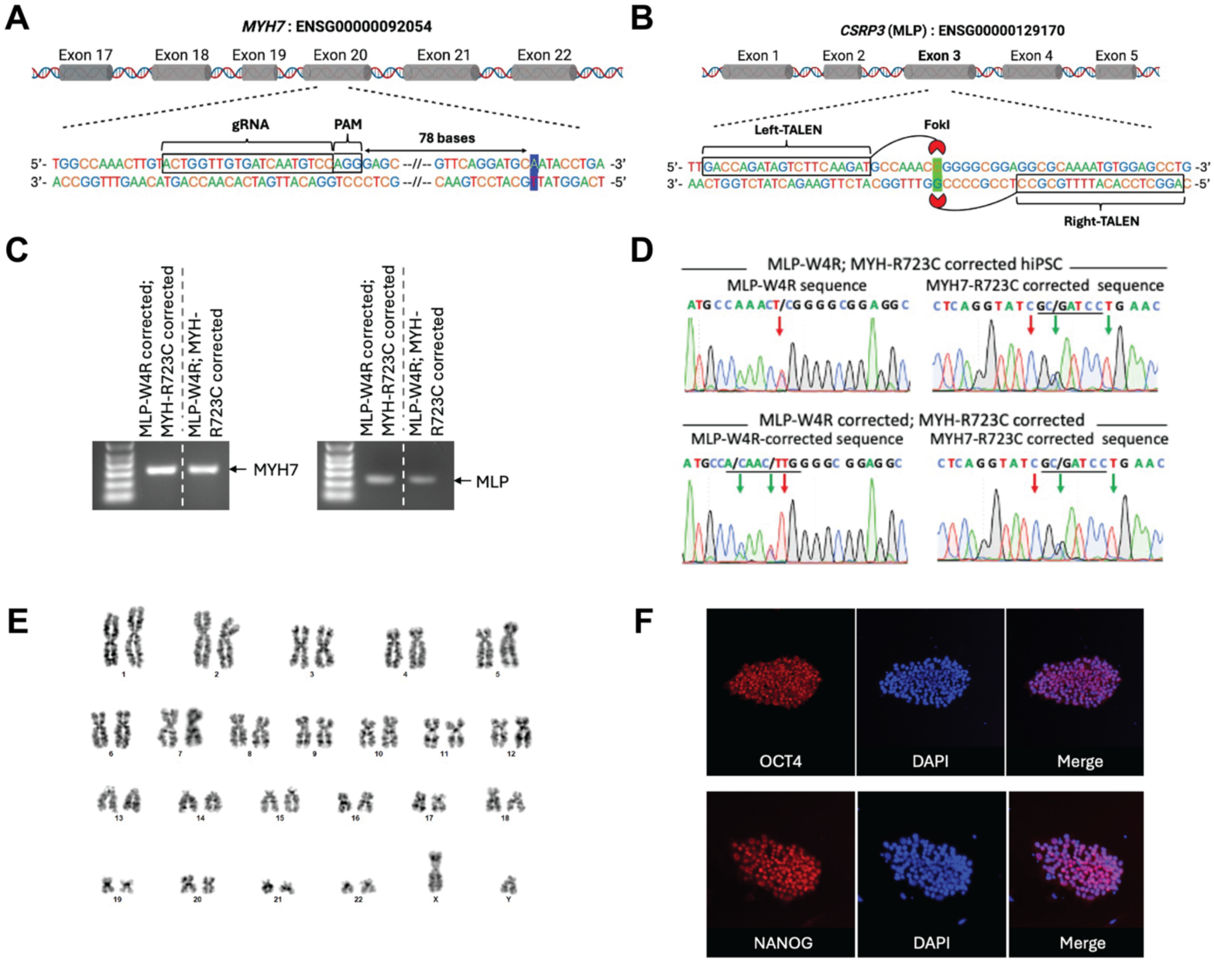


**Figure S2. Related to Figure 4. Generation and verification of isogenic control of Isogenic MLP-W4R iPSCs.**

**A**. Schematic of CRISPR/Cas9-mediated correction of the heterozygous MYH7-R723C missense mutation in exon 20. The guide RNA (gRNA) sequence (minus strand) and its protospacer adjacent motif (PAM) are highlighted. The target mutation (C to T) is located 78 base pairs downstream of the PAM sequence in MLP-W4R;MYH7-R723C proband iPSCs.

**B.** Schematic of TALEN-mediated correction of the MLP-W4R mutation in exon 3 of the CSRP3 gene. The left and right TALEN DNA-binding domains, fused with the FokI cleavage domain, flank the target mutation (T to C).

**C-D.** Agarose gel images and Sanger sequencing results confirm the MYH7-R723C and MLP-W4R corrections in IsoMLP-W4R iPSCs and IsoCtrl iPSCs (MYH7-R723C; MLP-W4R both corrected; isogenic control iPSCs).

**E.** Karyotype analysis of the MYH7-R723C;MLP-W4R doubly corrected IsoCtrl-iPSCs clone showing a normal karyotype.

**F.** Immunofluorescence staining of OCT4 and NANOG to assess pluripotency in IsoCtrl-iPSCs. Note that IsoMLP-W4R iPSC line was characterized to have normal karyotyping and pluripotency and reported in our previously reported study[1].

**Figure S3**

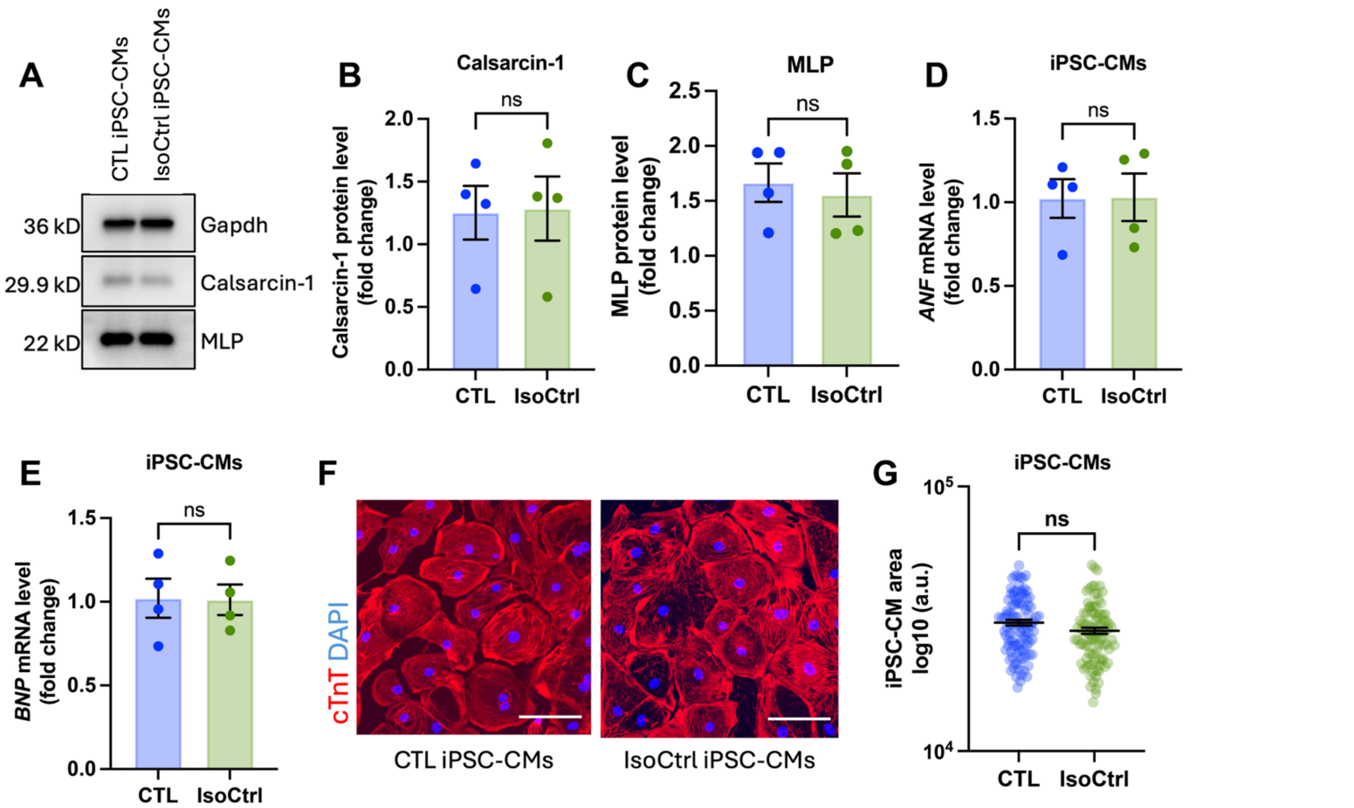


**Figure S3. Related to Figure 4. Cardiomyocytes derived from healthy donor CTL iPSC and isogenic control iPSC exhibited comparable characteristics.**

**A-C.** Calsarcin-1 and muscle LIM protein expression levels of cardiomyocytes derived from healthy donor CTL iPSC (CTL iPSC-CMs) and isogenic control iPSC (IsoCtrl iPSC-CMs), *n*=4 independent batches of cardiac differentiation, ns: not significant (2-tailed unpaired Student's t-test).

**D-E.** The mRNA expression levels of atrial natriuretic factor (*ANF*) and brain natriuretic peptide *(BNP)* in CTL iPSC-CMs and IsoCtrl iPSC-CMs, *n*=4 independent batches of cardiac differentiation, ns: not significant (2-tailed unpaired Student's t-test).

**F-G**. Representative images of cardiac troponin T (cTnT) immunofluorescence staining and cell size in CTL iPSC-CMs and IsoCtrl iPSC-CMs, *n*=4 independent batches of cardiac differentiation, ns: not significant (2-tailed unpaired Mann-Whitney U test). All data are presented as mean ± SEM.

**Reference:**

1 Riaz, M. *et al.* Muscle LIM Protein Force-Sensing Mediates Sarcomeric Biomechanical Signaling in Human Familial Hypertrophic Cardiomyopathy. *Circulation* **145**, 1238-1253 (2022). https://doi.org:10.1161/CIRCULATIONAHA.121.056265
